# Imperfect hairpins formed by CTG trinucleotide repeats are heteroduplex DNA molecules in living cells

**DOI:** 10.1101/2024.06.04.597286

**Authors:** David Viterbo, Pauline Dupaigne, Pierre Alexandre Kaminski, François Dejardin, Stéphane Petres, Eric Le Cam, Guy-Franck Richard

## Abstract

There are over fifty microsatellite expansion disorders, often neurodegenerative pathologies, due to large expansions of tandem repeat sequences. These microsatellites vary in motif composition, size and copy number in the genome. The ability of these repeats to form secondary structures *in vitro* is strongly suspected to trigger the expansion process and therefore, the pathology. However, there is no formal proof that similar structures are formed *in vivo*, in living and replicating cells.

We are focused on the CTG trinucleotide repeat whose expansion is responsible for myotonic dystrophy type 1 (DM1 or Steinert disease), affecting 1 in 8,000 people worldwide.

A series of llama nanobodies (V_H_H) were selected to bind to the secondary structure formed by a synthetic imperfect CTG hairpin. Yeast one-hybrid was used to identify V_H_H that were able to activate a reporter gene *in vivo* . One of them was shown to be very specific since it binds a CTG hairpin but not a CAG hairpin or a GC-rich sequence. This nanobody also bind to long CTG trinucleotide repeats carried by plasmidic or genomic DNAs, proportionally to the number of repeats, suggesting that longer repeats tend to form more frequently imperfect hairpins. Finally, the V_H_H was bound to a protein G column and used to selectively enrich structure-containing DNA molecules, that were further observed by electron microscopy. They exhibit different kinds of structures in which only one strand is folded into an imperfect hairpin.

Our results unambiguously show that alternative DNA structures are transiently formed by CTG trinucleotide repeats in living bacterial and yeast cells and suggest that imperfect CTG hairpins mainly exist *in vivo* mainly as heteroduplex DNA molecules.

## Introduction

Microsatellite expansions are responsible for more than two dozen neurological or developmental disorders. Among them, trinucleotide repeats are the most common microsatellite expanded. Since the discovery that CAG, CTG and CGG repeats were able to form secondary structures in defined conditions of salt, temperature and pH (Gacy et al. 1995; Mitas et al. 1995b, a; Yu and Mitas 1995; Mitas 1997; Yu et al. 1997; Suen et al. 1999), it was proposed that these structures may be the initial event triggering the expansion mechanism (McMurray 1999). Despite these initial observations, there is a lack of formal evidence showing that the same exact structures are formed in living and replicating cells.

Genetics studies in *Escherichia coli* and *Saccharomyces cerevisiae* show that CAG/CTG trinucleotide repeats are more unstable when the CTG-containing strand is the lagging strand template, suggesting that instability is linked to the stability of secondary structures formed on this strand (Kang et al. 1995; Freudenreich et al. 1997; Viterbo et al. 2016). Transient replication fork pausing occurs at long CAG/CTG trinucleotide repeats cloned in a plasmid (Samadashwily et al. 1997; Pelletier et al. 2003) or integrated in a yeast chromosome (Viterbo et al. 2016). In HeLa cells, a zinc-finger nuclease (ZFN) made of one arm binding to the (AGC)_3_ sequence (ZFN_CAG_) and one arm binding to the (GCT)_3_ sequence (ZFN_CTG_) was used to induce a double-strand break into a CAG/CTG trinucleotide repeat, triggering frequent contractions and expansions of the repeat tract. Expression of only one arm (ZFN_CAG_ or ZFN_CTG_) induced partial contractions (Liu et al. 2010). This was interpreted as the formation of a CAG (or CTG) DNA hairpin bound by a homodimeric ZFN_CAG_ or ZFN_CTG_ arm. However, CAG and CTG hairpins cannot fold as canonical Watson-Crick double-stranded DNA helices, since only G-C bonds are made (Figure 1A). Therefore, the possibility that a ZFN arm could bind such a structure may be debated.

**Figure 1:**
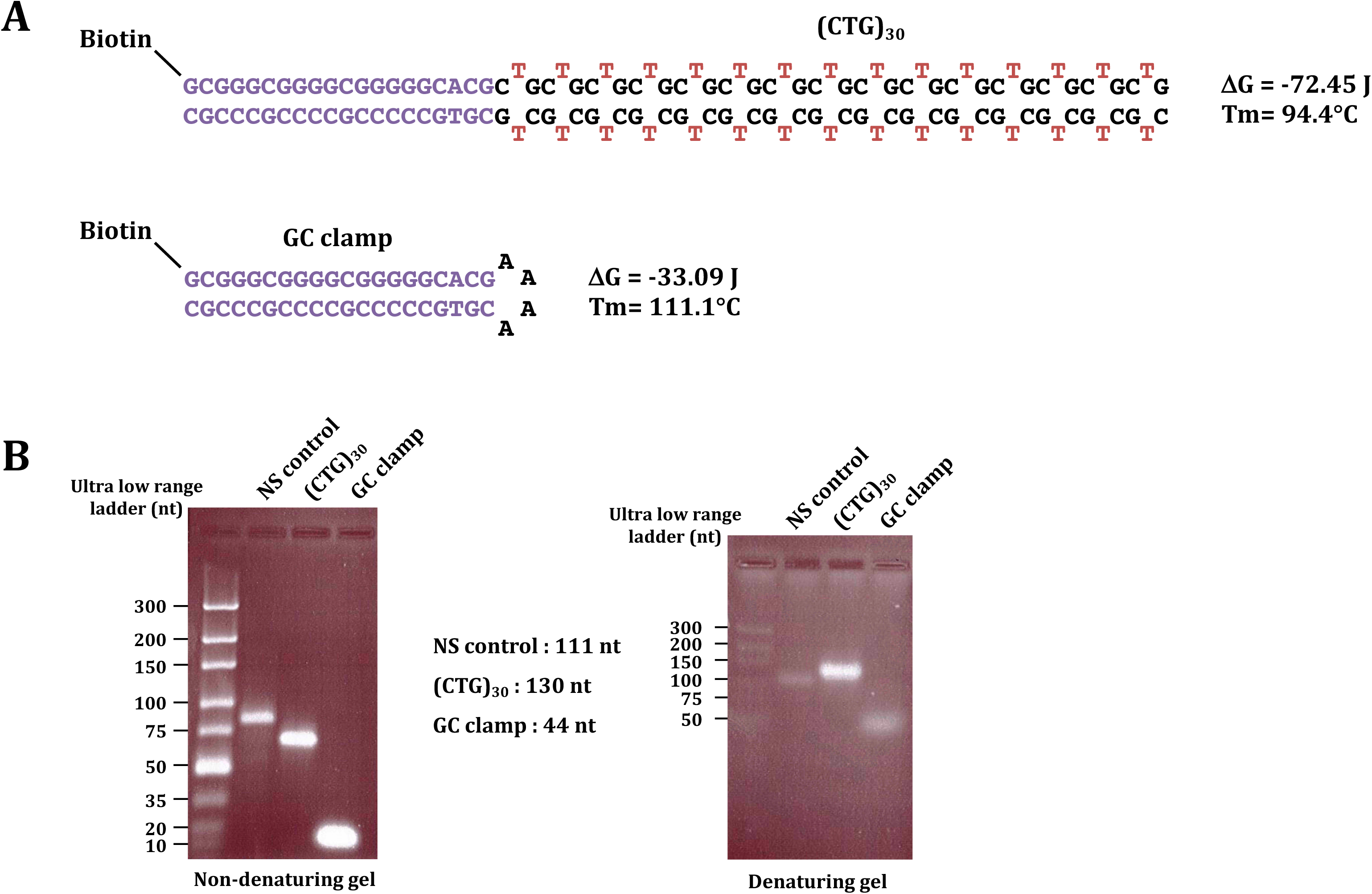
Design of the synthetic CTG hairpin. **A:** A 130 nt DNA oligonucleotide was designed in such a way that its ends are complementary and form a “GC clamp”, that tolerate formation of an imperfect hairpin in only one conformation. The GC clamp itself is synthesized to be used as a negative control in all experiments. Both oligonucleotides are linked to a biotin at their 5’ ends. Structures, free energy and Tm of both molecules were calculated using mFold version 3.6 (Zuker 2003). **B:** Migration of the (CTG)_30_ oligonucleotide, the GC clamp and a non structured (NS) 111 nt long oligonucleotide on, non-denaturing (left) and denaturing (right) agarose gels (see Materials and Methods for details). On the non-denaturing gel (3% Metaphor agarose (Lonza), TBE 1X), both the CTG hairpin and the GC clamp migrate as predicted for double-stranded DNA of these expected molecular weights. Note that the ethidium bromide signal is much stronger than for the NS oligonucleotide. On the denaturing gel (3% Metaphor agarose (Lonza), 50 mM NaOH, 1 mM EDTA) all three species migrate at their expected molecular weight. The ethidium bromide signal is stronger for the CTG hairpin, suggesting the possibility that at least part of the oligonucleotide may be double-stranded. Amounts loaded on each gel are approximately : NS control oligonucleotide: 20 pmole, CTG hairpin: 5 pmole, GC clamp: 28 pmole.

A monoclonal antibody (2D3) was raised against a cruciform (Steinmetzer et al. 1995) and used to immunoprecipitate CAG/CTG slipped-stranded DNA structures at the myotonic dystrophy type 1 (DM1) locus in several patient tissues (Axford et al. 2013a). More recently, naphthyridine-azaquinolone (NA), a synthetic molecule, was found to bind selectively *in vitro* to long CAG hairpins (but not to CTG hairpins), inhibiting their repair by HeLa cell extracts and triggering contractions of a CAG/CTG repeat tract in a mice model of Huntington’s disease (Nakamori et al. 2020).

All these experiments were taken as evidence that some kind of secondary structures were formed by CAG/CTG trinucleotide repeats *in vivo* . However, their precise nature, their presence in replicating or non-replicating DNA, their stability over time is still questionable (Poggi and Richard 2021). In order to directly and irrefutably address these questions, we use a phage display selection screen to identify several single chain nanobodies (V_H_H) specifically recognizing an imperfect CTG hairpin. These V_H_H were expressed in yeast one-hybrid assays and showed specific binding to CAG/CTG trinucleotide repeats *in vivo* . Out of the four that were purified as proteins, one showed a very specific binding to a CTG hairpin and not to a CAG hairpin or to a GC-rich sequence. Additional experiments show that this antibody recognizes CTG hairpins in plasmidic DNA but also in genomic DNA extracted from yeast and containing long CTG trinucleotide repeats from a human DM1 patient. Finally, the nanobody was bound to a protein G column and used to capture CTG hairpin-containing plasmids. Observation of these captured molecules by electron microscopy show that they exhibit a variety of length and shapes but suggest that most of them are found in cells as heteroduplex DNA molecules in which only on strand is folded into an imperfect hairpin.

## Results

### CTG hairpin design and phage display screen

We started by designing a synthetic piece of single-stranded DNA, containing 30 CTG triplets, ending with two short (20 nt) complementary single-strand DNA sequences forming a GC clamp (Figure 1B) (Sheffield et al. 1989). The rationale was to force the folding of the synthetic oligonucleotide into one single conformation, avoiding generating multiple slipped-stranded DNA species. A biotin was attached to its 5’ end for the phage display screen, and the GC clamp alone was also synthesized to deplete the library from non-specific binders. Denaturing and non-denaturing gels were run to verify that both oligonucleotides migrate as expected. On a non-denaturing gel, the 130 nt CTG hairpin migrates at a molecular weight corresponding to 63 bp, whereas the 44 nt GC clamp alone migrates as an 18 bp species (Figure 1B, left). A supposedly non-structured 111 nt control oligonucleotide migrated as an 82 bp species, suggesting that a portion of it is double-stranded. Only one DNA species could be detected in each lane of the non-denaturing gel. When the same DNA species were run on an alkaline denaturing gel, all three of them migrated at their expected molecular weight (Figure 1B, right). This confirmed that both the CTG hairpin and the CTG clamp are folded into the expected secondary structures.

Using the CTG hairpin as a bait, three successive rounds of phage display were performed (Materials & Methods). Progressive enrichment was observed by the ratio output/input and by ELISA assay (Supplemental Figure S1). After the third round, 83 out of 90 phages tested by ELISA assay were positive for binding to the CTG hairpin. Sequencing of the positive clones revealed 27 different V_H_H sequences, some of them being found several times. Eleven of them were selected, based on ELISA results and on their redundancy (Table 1).

### One-hybrid assays

All of them were cloned in a one-hybrid yeast plasmid (pP9), in frame with the *GAL4* binding domain (GBD), under the control of the strong constitutive *ADH1* promoter. The 11 plasmids were each transformed into a *MAT*a strain. A *MAT*α strain containing a (CTG)_99_ trinucleotide repeat upstream the *HIS3* gene under the control of a minimal promoter was built (Materials & Methods). If the V_H_H-GBD fusion binds to a hairpin formed by the CTG repeat, it will activate transcription of the downstream reporter gene and cells will be [His+] (Figure 2A). Each of the 11 haploid *MAT*a strain was crossed to the *MAT*α reporter strain, diploids were selected and spotted on synthetic medium lacking histidine (SC -His), at two different temperatures (Figure 2B). As expected, cells grew faster at 30°C as compared to 20°C, but the general trend, as compared to the control strain with the pP9 alone was the same. Each V_H_H-GBD fusion was given a score according to growth on SC -His. Yeast cells containing the C09 and the D11 fusion did not grow and were given negative scores as compared to the control. Yeast cells containing the G05 and F08 fusions grew as the control, whereas seven V_H_H-GBD fusions showed a better growth than the control on Sc -His. We concluded that these V_H_H were able to bind the (CTG)_99_ repeat in yeast cells, suggesting that it was able to form, at least transiently, secondary structures resembling the hairpin used in the phage display screen.

**Figure 2:**
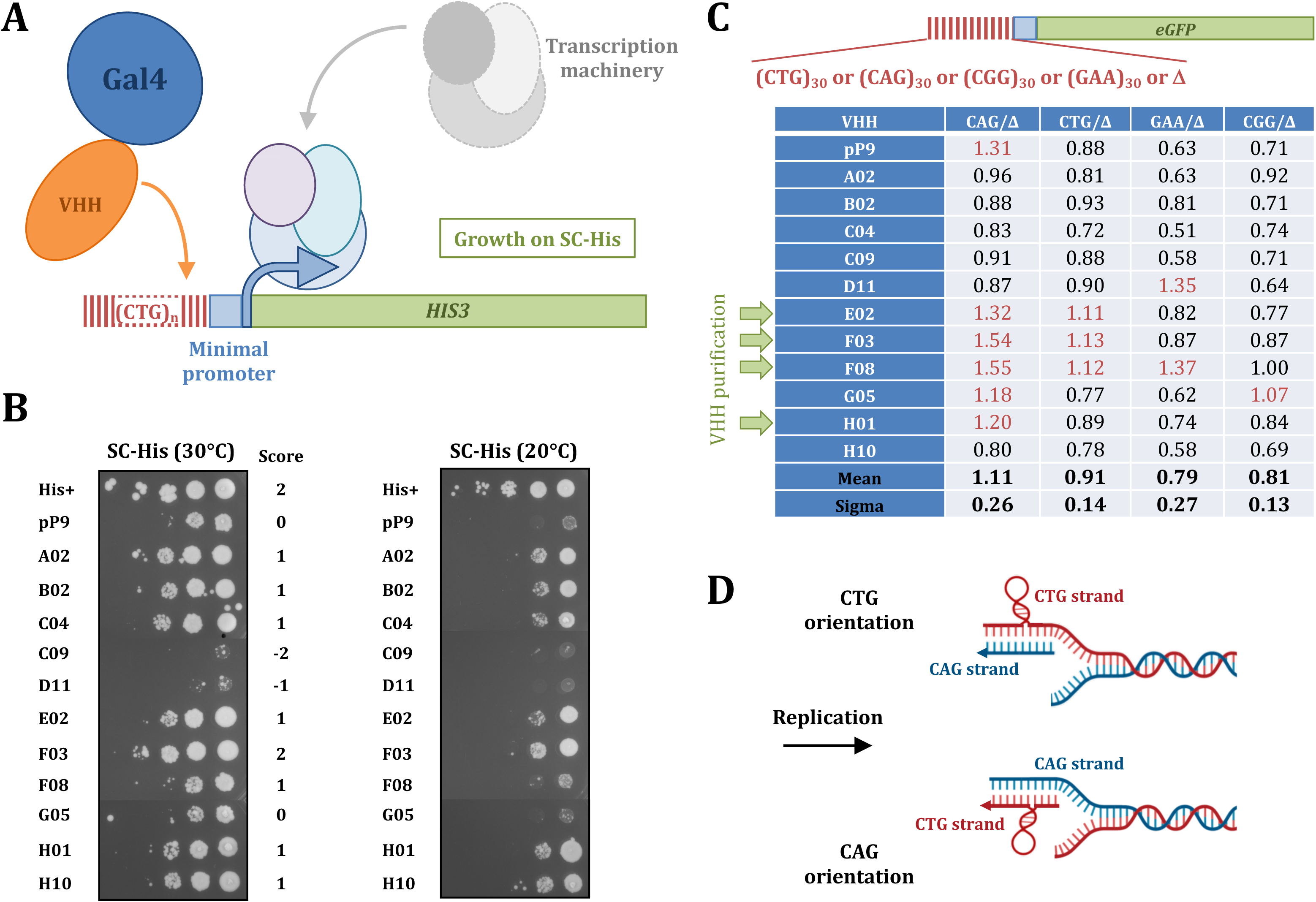
One-hybrid experiments. **A:** The *HIS3* gene under the control of a minimal promoter fused to a CTG trinucleotide repeat was integrated in the yeast genome. The 11 V_H_H fused to the Gal4p activating domain were expressed in this yeast strain. If the fusion protein binds to the CTG repeat it will activate transcription of the *HIS3* gene and cells will grow on minimal medium without histidine (SC-His). **B:** Spot test on SC-His. A [His+] strain was used as positive control and the plasmid containing the *GAL4* activating sequence but no V_H_H (pP9) allowed to determine the baseline of growth on SC-His. The score represents the difference of growth between each V_H_H-Gal4 fusion and the pP9 vector. Note than some V_H_H obtained a negative score, due to a worse growth than the pP9 control. Yeast plates were incubated at two different temperatures, 30°c which is the physiological temperature for budding yeast and 20°C. As expected, growth was slower at lower temperature, but it did not change the general outcome. **C:** One-hybrid results using a *GFP* reporter gene under the control of the minimal *HIS3* promoter. The promoter was fused to four different microsatellites: (CTG)_30_, (CAG)_30_, (GAA)_30_ or (CGG)_30_. An isogenic construct in which no microsatellite was present was used as a control of baseline fluorescence (Δ). The table shows the average ratio of the GFP fluorescence of each repeat tract divided by the baseline fluorescence (Δ), at the two time points after alpha factor release (see text and Supplemental Table S1 for details). Values above 1 indicate a higher fluorescence with a repeat tract than with no repeat (Δ) and are shown in red. The four green arrows point to V_H_H that gave the best one-hybrid results both with the *HIS3* reporter and the *GFP* reporter and were produced as proteins. **D**: Cartoons depicting the position of the CTG hairpin on the lagging strand template (CTG orientation) or the Okazaki fragment (CAG orientation), depending on the orientation of the CAG/CTG repeat tract (see text for details).

In order to verify the binding specificity, we designed a similar one-hybrid screen, in which the *HIS3* reporter was replaced by the enhanced *GFP* gene and the (CTG)_99_ was replaced by a synthetic (CTG)_30_, (CAG)_30_, (GAA)_30_ or (CGG)_30_ trinucleotide repeat. These four synthetic genes were integrated at the *ARG2* locus, as well as a control construct without any repeat. Yeast cells were first arrested with alpha factor, then released in fresh medium without alpha factor and collected at the beginning of the experiment, after 120 minutes and 180 minutes, covering the entirety of the cell cycle in this medium. The number of GFP-positive cells was determined at each time point and the ratio over cells without any repeat was calculated (Supplemental Table S1). No general trend of increase or decrease in GFP-positive cells over time was visible. We therefore used the average of the three time points to compare the results (Figure 2C). The average GFP-positive values for the (CAG)_30_ are overall above the (GAA)_30_ and (CGG)_30_values (t-test p-value= 1.2 x 10^-7^ for GAA and 3.4 x 10^-13^ for CGG). It is also the case for (CTG)_30_ (t-test p-value= 9.6 x 10^-4^ for GAA and 4.3 x 10^-7^ for CGG). The same holds true when ratios between microsatellite-containing strains and the strain containing no repeat (Δ in Figure 2C) are considered.

It was expected that (CAG)_30_ and (CTG)_30_ repeats behaved similarly, since a CTG hairpin may form on each of these double-stranded DNA sequences. However, (CAG)_30_ figures are significantly above (CTG)_30_, showing that CTG hairpins form more readily in this orientation (t-test p-value= 2.1 x 10^-4^).

Interestingly, the (GAA)_30_ and (CGG)_30_ are less efficient than the Δ strain to activate the GFP reporter (t-test p-value= 1.1 x 10^-5^ for GAA and 1.5 x 10^-11^ for CGG). This suggests that these two repeats inhibit non-specific Gal4 activation of the reporter gene.

In the strains containing (CTG)_30_ or (CAG)_30_ repeats, three V_H_H and five V_H_H respectively showed ratios above 1. Indeed, the three V_H_H that showed ratios above 1 in the (CTG)_30_ strain, were also positive in the (CAG)_30_ strain (E02, F03 and F08, Figure 2C). Note that, unexpectedly, F08 also shows a GFP-positive signal with the (GAA)_30_ strain. This suggests that this nanobody may recognize an epitope that is common to both CTG and GAA structures. Nevertheless, these three V_H_H, as well as H01, which was also a good candidate in the Histidine one-hybrid assay, were purified as proteins fused to a rabbit Fc domain (see Materials & Methods).

### In vitro assays

The CTG hairpin and the GC clamp were immobilized on a nylon membrane and hybridization with each of the four purified V_H_H was performed (see Materials & Methods). Only H01 gave a strong positive signal with the hairpin, in these conditions (Figure 3A, top). This V_H_H was therefore used in all the subsequent *in vitro* experiments. It was shown to be very specific of a CTG hairpin, since no signal was detected when using a CAG hairpin of the same length or the GC clamp itself (Figure 3A, bottom). Subsequently, increasing amounts of plasmids containing either 98 or 255 CTG triplets were hybridized with the same V_H_H. Plasmids pRW3216 and pRW3222 are pUC19 derivatives in which a (CTG)_98_ or a (CTG)_255_ triplet repeat from a patient affected by DM1 was cloned (Richard et al. 2000; Fabre et al. 2002). The signal was much stronger with the plasmid containing 255 CTG triplets (Figure 3B). We performed the same experiment with or without preliminary heat denaturation followed by fast renaturation of the plasmids. The rationale was that denaturation/renaturation should artificially increase the number of secondary structures carried by plasmids. The signal was indeed much stronger when plasmids were heat denatured. No signal was detected in any condition with the pUC19 itself (Figure 3C). Finally, we hybridized total yeast genomic DNAs in which different lengths of chromosome-borne CTG trinucleotide repeats were integrated (Viterbo et al. 2016). The signal was stronger with DNAs that contained CTG triplets than with the wild-type strain (Figure 3D). It increased to a length of 100 triplets before decreasing above that, suggesting that alternative structures may form that are not anymore recognized by the V_H_H. As a positive control, human DNA containing a very large CTG repeat expansion at the *DMPK* locus (Arandel et al. 2017) exhibited a much stronger signal. We concluded from this series of experiments, that the H01 V_H_H was able to specifically bind to a CTG hairpin *in vitro*, either on a single-stranded oligonucleotide, on a plasmid or on total genomic DNA.

**Figure 3:**
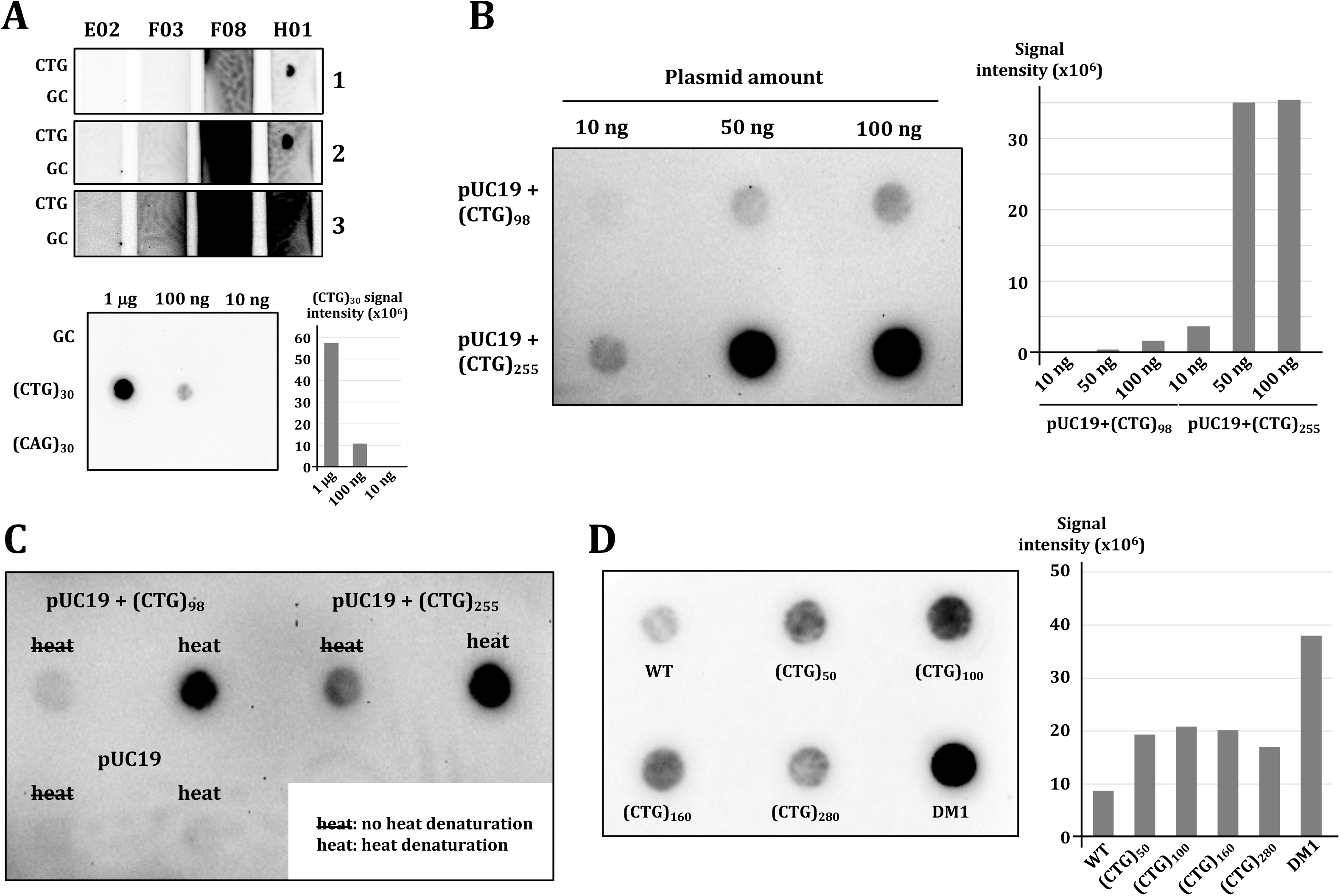
Dot blots with purified V. _H_**H proteins.** A: Top: Nylon membranes on which 100 ng of the synthetic CTG hairpin (CTG) and of the GC clamp (GC) were spotted were incubated with one of the four V_H_H (E02, F03, F08 and H01). Three pictures were taken at different exposure times (labeled 1, 2 or 3). Only H01 gave a significant positive signal with the CTG hairpin and no signal with the GC clamp. Bottom: H01 was incubated with a membrane on which different amounts of the CTG hairpin (CTG), the GC clamp (GC) or a synthetic CAG hairpin (CAG) were spotted. A positive signal was detected only with the CTG hairpin, and its quantification is shown to the right. **B:** The V_H_H H01 reveals the presence of CTG hairpins on plasmidic DNA containing 98 or 255 CTG triplets. Signal quantifications are shown to the right. **C:** Heat denaturation of plasmidic DNA creates additional CTG hairpins that are detected by the V_H_H H01. No signal is visible with the same plasmid containing no repeat tract. **D:** Yeast total genomic DNA containing CTG trinucleotide repeats of different lengths (50, 100, 160 or 280 triplets), as well as human total genomic DNA of a DM1 patient contain different amounts of CTG hairpins, as revealed by the V_H_H H01. Signal quantifications show that genomic DNA containing 160 and 280 triplets exhibit a lower signal than genomic DNA containing shorter repeat lengths. The DM1 genome contains 2600 CTG triplets at the *DMPK* locus (Arandel et al. 2017).

### Purification and visualization of secondary structures by electron microscopy

In a subsequent set of experiments, the V_H_H was bound to a HiTrap protein G column by its Fc domain (see Materials & Methods) and used as a bait to capture structure-containing plasmids, freshly extracted from *E. coli* . Plasmids pRW3222 containing 255 CTG triplets and the original pUC19 were used. In a first experiment, native pRW3222 extracted from *Escherichia coli* was loaded on a HiTrap column in which the H01 V_H_H was bound. After increasing salt concentration from 50 mM to 200 mM, the eluted fractions were enriched in plasmid, proving that some of the native plasmids were bound to the column (Supplemental Figure S2). As a control, when the same plasmid was loaded on a column that did not contain the V_H_H, no elution was observed after increasing salt concentration.

We subsequently performed the same experiment with both plasmids linearized with *Ssp*I, a restriction enzyme that cuts only once in pUC19, 646 bp before the CTG repeat tract. Following digestion, both plasmids were heat denatured, before being loaded on a column. The reasoning was that linearized and denatured plasmids should form secondary structures more readily that native plasmids. pRW3222 and pUC19 were identically treated and each of them was loaded on a protein G column. A sample of each collected fraction was loaded on an agarose gel, transferred to a nylon membrane and hybridized with a pUC19-specific probe. Signal quantification showed that 4.9% of the pRW3222 was eluted in the high salt fractions, compared to only 0.3% of the pUC19 (Figure 4). Eluted pRW3222 plasmids were afterwards analyzed by electron microscopy (see Materials & Methods). Out of 41 molecules, 18 exhibited a single-stranded region adopting different conformations, hairpin-like or more complex structures (Supplemental Figure S3). The single-stranded regions are located on the average at 23% of plasmid length, which is the location of the CTG trinucleotide repeat (Supplemental Table S4). This is the demonstration that despite its GC-rich content, this region is more prone to heat denaturation than the rest of the plasmid, suggesting that it adopts a non-canonical Watson-Crick conformation, and that it renatures in different non-B DNA conformations that are bound by the anti-CTG V_H_H.

**Figure 4:**
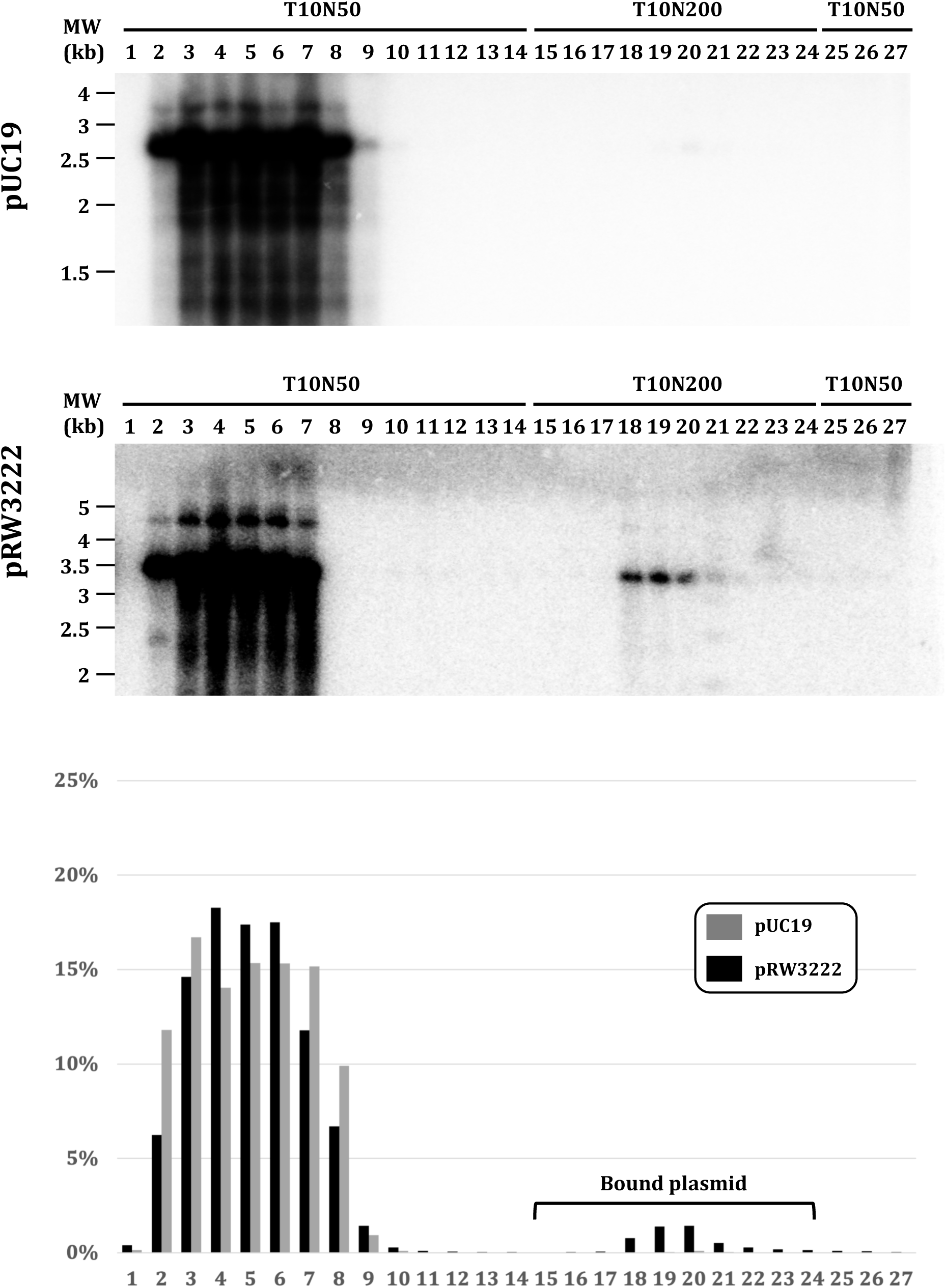
Enrichment of denatured CTG-containing DNA on a protein G-V_H_H column. Linearized and denatured pUC19 (20 μg) or pRW3222 (20 or 100 μg) were loaded on a HiTrap protein G column on which the H01 V_H_H-Fc was bound (flow rate= 1 mL/min). A sample of each collected 1 mL fraction was loaded on an agarose gel, which was subsequently transferred to a nylon membrane and radioactively hybridized with a pUC19 probe. The pRW3222 is eluted in high salt fractions (15 to 24), whereas the pUC19 is barely detectable in the same fractions (see text for details). Quantifications were performed on a Fujifilm FLA-9000 phosphor screen reader. T10N50: 10 mM Tris pH 8.0, 50 mM NaCl; T10N200: 10 mM Tris pH 8.0, 200 mM NaCl.

In order to determine more precisely the kind of structures in which CTG hairpins are folded *in vivo*, the same experiment was subsequently conducted on native pRW3222. Plasmidic pRW3222 DNA was extracted from *E. coli* and directly loaded on a protein G column, in the same conditions as before. Samples from collected fractions were loaded on an agarose gel, transferred to a nylon membrane and hybridized with the pUC19 probe. Two bands were visible, respectively corresponding to supercoiled and relaxed plasmids. Quantification of the signal showed that 0.9% of the plasmid was eluted when increasing salt concentration, roughly in the same proportion for the two molecular species (Figure 5). Eluted plasmids were afterwards analyzed by electron microscopy, as previously. Out of 58 plasmids, 35 contained non-canonical Watson-Crick conformations (60%), 10 of them containing two different structures (Supplemental Table S5). Lengths of the structures ranged from 4 to 82 nucleotides. Given that the CTG tract is 765 bp long (3 x 255 CTG triplets) the observed structures do not cover the entirety of the repeat tract. Altogether, 45 structures were observed, whose mean length was 30 bp (median length= 27 bp). All the pRW3222 plasmids studied are shown in Supplemental Figure S4 (plasmids #1 to #58).

**Figure 5:**
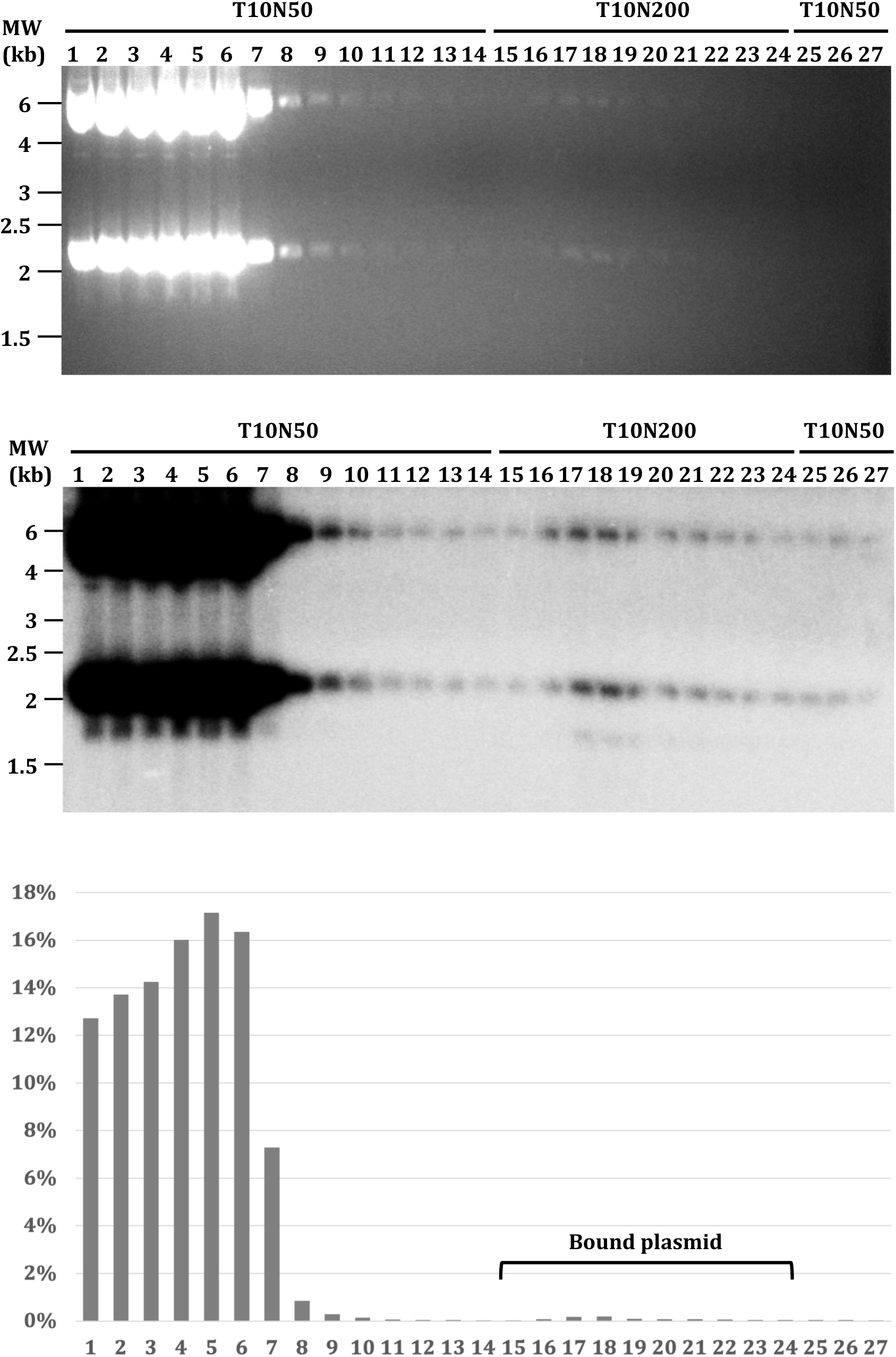
Enrichment of native CTG-containing DNA on a protein G-V_H_H column. Native pRW3222 (500 μg) were loaded on a HiTrap protein G column on which the H01 V_H_H-Fc was bound (flow rate= 0.5 mL/min). **Top** : Fractions collected from the FPLC were loaded on a non-denaturing agarose gel (Top), run at 4V/cm in 1X TBE buffer in the presence of 0.02 μg/mL ethidium bromide. The gel was transferred and hybridized with radioactively labeled pUC19 DNA. Supercoiled and relaxed forms of the pRW3222 plasmid are visible as discrete bands. The total signals detected in each lane are quantified on a Fujifilm FLA-9000, as previously described (Viterbo et al. 2018). **Bottom** : Quantification results are given in arbitrary units. The amount of structured plasmid eluted from the V_H_H after increasing salt concentration (T10N200) corresponds to 0.9% of the total plasmid loaded on the column.

All the observed secondary structures were classified according to their shape. Long hairpins were called type A; short hairpins were called type B and all other shapes (bent hairpins, eye of a needle-like hairpins, thicker than usual hairpins, etc.) were called type C. Type A are the most frequent and represent almost half of the observed structures, followed by type B and C in roughly the same proportions (Figure 6). No structure like a cruciform could be observed, strongly suggesting that when a CTG hairpin is formed on one strand a CAG hairpin is not formed on the opposite strand, or if it is formed it is not visible by electron microscopy. It is rather surprising that in 10 cases out of 58 molecules, two structures were seen at two different places, suggesting than non-CTG sequences on the plasmid may also sometimes form secondary structures.

**Figure 6:**
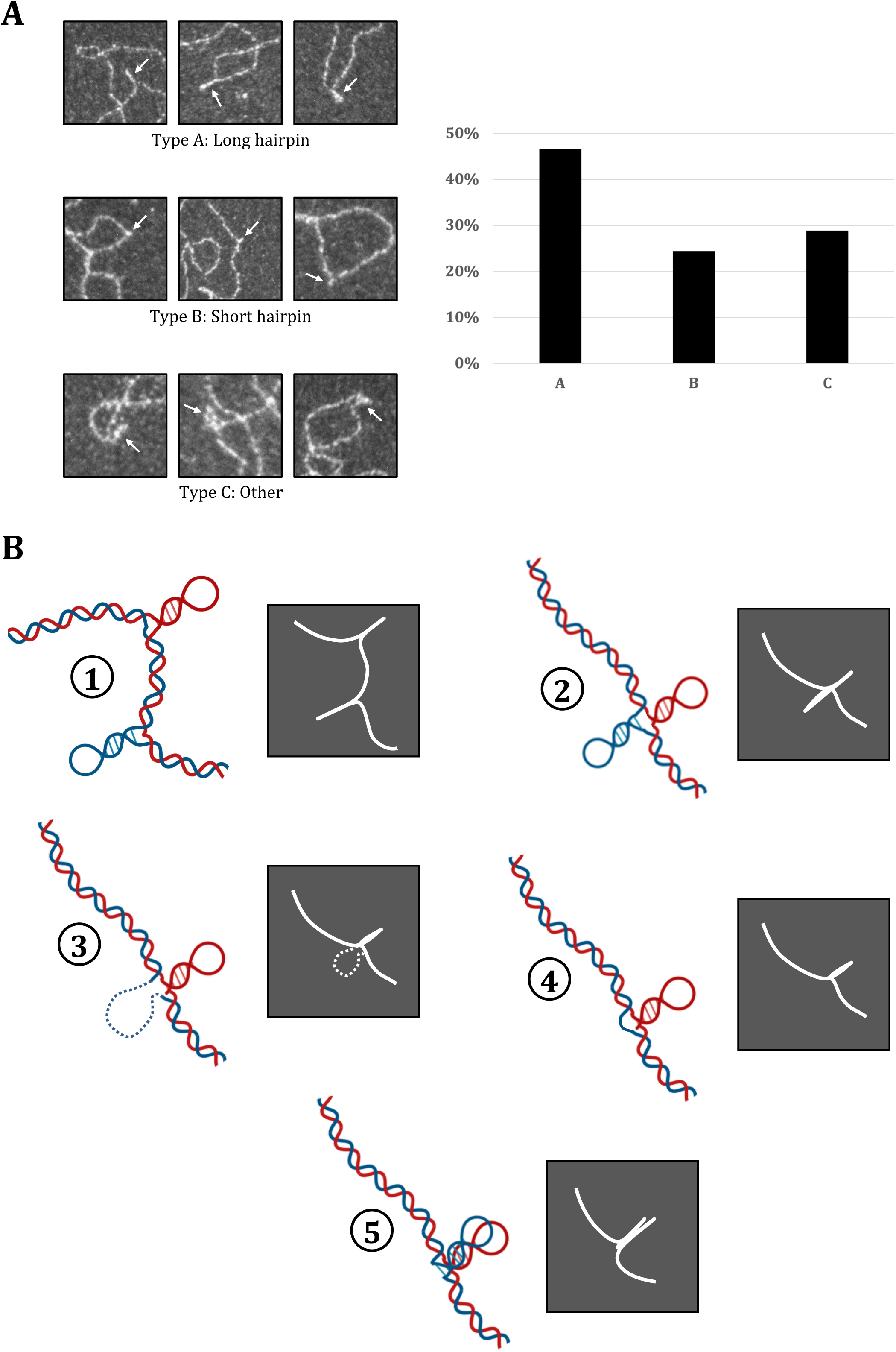
Different types of secondary structures visible par electron microscopy. **A**: Some examples of the three types of structures observed, with their proportions in the pRW3222. **B** : Cartoons depicting the different kinds of DNA structures that could be expected and their corresponding image by EM (see text for details).

## Discussion

In the present work, we have identified several nanobodies directed against an imperfect CTG trinucleotide repeat hairpin, that are able to activate transcription in two different reporter systems in *S. cerevisiae*, proving that such structures are formed in dividing yeast cells. Previous works using genetics and molecular biology showed that the presence of a CTG trinucleotide repeat integrated in a plasmid or a yeast chromosome altered replication, repair and, more generally, DNA metabolism (reviewed in Poggi and Richard 2021). However, until now, there was no formal demonstration that these repeats were actually able to form the same imperfect hairpins *in vivo* as *in vitro* . Our results prove that this is the case.

The H01 V_H_H was shown to be active as an intrabody in *S. cerevisiae* cells (Figure 2). Among the four nanobodies tested on a nylon membrane, it was the only one to give a positive signal on an immobilized CTG hairpin in the conditions of our experiment (Figure 3A). Interestingly, it was also the best candidate in the ELISA test and was found 15 times during the screening process (Table 1). Finally, we showed that it selectively binds to plasmids extracted from *E. coli* and exhibiting different kinds of hairpins as demonstrated by electron microscopy (Figures 4 to6). All these properties make it the best nanobody to be used in future experiments with CTG trinucleotide repeats.

The H01 V_H_H binds to a CTG hairpin but not to a CAG hairpin (Figure 3). This shows that, although DNA is double-stranded, positive signals detected in one-hybrid experiments are due to the formation of CTG hairpins only. Interestingly, in the e*GFP* one-hybrid assay, the signal is significantly higher in the CAG than in the CTG orientation (Figure 2C). Position of the closest active replication origin, upstream the reporter gene, shows that in the CAG orientation, the CTG sequence is on the Okazaki fragment, whereas in the CTG orientation, it is on the lagging-strand template. Previous experiments in bacteria and budding yeast have shown that CAG/CTG triplet repeats are more unstable in the CTG than in the CAG orientation (Kang et al. 1995; Freudenreich et al. 1997; Viterbo et al. 2016). It was therefore proposed that CTG hairpins form more readily on the lagging-strand template than on Okazaki fragments. Our results suggest the opposite, since the signal indicative of the presence of a CTG hairpin is slightly stronger in the CAG orientation (Figure 2D). Note that we cannot totally exclude than CTG hairpins also form on the leading-strand template, which carries the same sequence as Okazaki fragments, but previous experiments using *rad27* mutants in yeast strongly argue in favor of a replication defect of CAG/CTG repeats on the lagging strand rather than on the leading strand (Freudenreich et al. 1998; Schweitzer and Livingston 1998; Richard et al. 1999).

### CTG trinucleotide repeats form an unexpected variety of structures in E. coli

Our experiments allowed, for the first time, to visualize by electron microscopy (EM) alternative DNA structures formed by CTG repeats cloned in a pUC19 plasmid extracted from *E. coli* . It was previously thought that such repeats could form slipped-strand DNA (Pearson et al. 2002; Axford et al. 2013b) or possibly cruciform-like structures like those formed by real palindromic sequences (Leach et al. 1997; Lobachev et al. 2000, 2002; Ait Saada et al. 2021). These putative structures are cartooned in Figure 6B (structures 1 and 2). However, we did not observe such structures by EM. There are three possible explanations for that.

Firstly, the hairpin is formed by the CTG strand, but there is no structure on the CAG strand (Figure 6B, structure 3). CAG-containing DNA is single-stranded and therefore hardly visible by EM. We may rule out this hypothesis since when the plasmid was formerly digested and denatured, single-stranded structures were visible as fuzzy lines linked to double-stranded DNA (Supplemental Figure S3). Nothing comparable was ever observed here.

Secondly, it was formerly shown that unrepaired CTG hairpins formed on one strand during replication and left unrepaired lead to heteroduplex molecules, detected as sectored colonies in yeast (Viterbo et al. 2016). Such DNA molecules carry one hairpin forming CTG repeat on one strand and a shorter CAG repeat on the other strand (Figure 6B, structure 4). This is compatible with our observations and strongly suggests that CTG hairpins do exist as heteroduplex DNA in replicating cells.

Finally, it is also possible that both CTG and CAG hairpins are formed very close to each other, separated by only one helix turn and therefore visible as a single hairpin (Figure 6B, structure 5). The form seen by EM would be slightly larger than a single hairpin. Some of the Type C structures could be such double hairpins (some examples are shown in figure 6B). This would also suggest that when a CTG hairpin is formed on one strand, a CAG hairpin is formed very close to it, at always the same distance, which cannot be completely ruled out but seems unlikely to frequently occur.

## Materials and Methods

### Plasmids

V_H_H sequences were PCR amplified with Phusion polymerase (NEBiolabs), with primers VHHup and VHHdown. They were cloned into the pP9 plasmid (Hybrigenics) digested by NcoI and NotI, by gap repair in the FYEF01-9C yeast strain, a MATa derivative of the FYAT01 strain (Richard et al. 1999) (Supplemental Table S3). pP9 contains the strong constitutive *ADH1* promoter driving the expression of the *GAL4* activation domain to which V_H_H were cloned in frame. The resulting plasmids pTRI136 to pTRI146 (Table 1) were sequenced (Eurofins) to verify the absence of any mutation and the integrity of each fusion gene.

To clone CTG repeats in the pB300 one-hybrid reporter plasmid (Hybrigenics), the repeats were gel purified from plasmid pRW3216 (Richard et al. 2000) digested with BamHI and PstI, and ligated to pB300 digested with BamHI and SalI, using complementary oligonucleotides pB300-Sal and pB300-Pst as adapters (Supplemental Table S2). The resulting pTRI148 plasmid was sequenced (Eurofins) and contains 99 uninterrupted CTG triplets, upstream the minimal *HIS3* promoter driving the expression of the *HIS3* gene.

One-hybrid GFP reporters were ordered as synthetic pieces of DNA from Thermo Fisher Scientific for CAG-GFP, CTG-GFP and GAA-GFP and from Eurofins for CGG-GFP. Each sequence contains exactly 30 triplets, flowed by the minimal *HIS3* promoter and the eGFP gene. Each synthetic gene was cloned at ApaI and BamHI in pTRI109 (Richard et al. 2003), to give plasmids pTRI149 (CAG), pTRI150 (CTG), pTRI151 (GAA) and pTRI152 (CGG), that contain flanking homologous regions with the *ARG2* gene. Plasmid pTRI150 was subsequently digested with BamHI and EagI, to remove the CTG repeat tract, treated with T4 DNA polymerase to blunt the extremities and religated to make pTRI153, which contains eGFP under the control of the *HIS3* minimal promoter and no repeat tract (“Δ” in FACS analyses).

### Yeast strains

Plasmid pTRI148 was linearized with HpaI in the *ADE2* gene and transformed in yeast strain FYEF01-7C, isogenic to FYAT01 carrying the *ade2*-opal mutation (Richard et al. 1999) (Supplemental Table S3). The resulting [Ade+] transformants were analyzed by Southern blot, to check correct integration at the *ADE2* locus as well as CTG repeat tract length. One clone, called GFY211, was selected and crossed to all FYEF01-9C transformants containing the pP9-V_H_H plasmids (pTRI136 to 146).

Plasmids pTRI149-152 containing eGFP constructs were digested by XhoI, transformed into FYEF01-7C and transformants were selected on SC-Trp. Since the plasmid is targeted to the *ARG2* gene, [Arg-] transformants were analyzed by Southern blot to check correct integration and repeat length. pTRI153 was digested with XhoI, transformed into FYBL2-7B, and [Trp+, Arg-] transformants were kept for further experiments. Strains GFY212 to 215 were subsequently switched to *MAT*a using pJH132, as previously described (Coïc et al. 2006). Ultra Low Range DNA ladder (Invitrogen) was used as molecular weight marker.

### Phage display

Synthetic CAG hairpin, CTG hairpin and GC clamp were ordered as PAGE Ultramer deoxynucleotides (IDT), linked to a 5’ biotin (Figure 1). A non-related antigen, GC Clamp-Biotin was used in an initial round of Phage Display to deplete the library from unspecific binders. Prior to the Phage Display selection, CTG Hairpin-Biotin and GC Clamp-Biotin were bound to Streptavidin Magnetic Beads (Dynabeads® M-280 Streptavidin, Life Technologies) with a 50 nM final concentration of biotinylated antigen for the first round and 10 nM final concentration of biotinylated antigen for the second and third rounds.

Three rounds of Phage Display selection were carried out using biotinylated CTG Hairpin. Hybrigenics’ synthetic hsd2Ab V_H_H library of 3.10^9^ clones (Moutel et al. 2016) was expressed at the surface of M13 phage. Hybrigenics’ Phage Display allowed to select V_H_Hs recognizing the non-adsorbed antigen in a native form. A total of 90 V_H_Hs were picked and analyzed after three rounds of Phage Display for binding to CTG hairpin, and 83 of these V_H_Hs were positive in non-adsorbed phage ELISA (Supplemental Figure S1). After sequencing of these 83 clones, 27 different clones were validated and positive in non-adsorbed phage ELISA. Among those, 11 clones were selected from the screen according to their criterion of redundancy, Elisa assay results and the sequencing results (Table 1). The 11 clones selected are Nali_PA00301_C09_woAmber, H01, A02, G05, E02, F03_woAmber, F08, H10, C04, D11 and B02. The two clones carrying Amber stop codons were corrected before being expressed in yeast.

### One-hybrid experiments

For the CTG-*HIS3* experiments, FYEF01-9C yeast cells containing the pP9-V_H_H plasmids were crossed to GFY211 containing the CTG-*HIS3* reporter gene. Diploids were selected on SC-Trp -Lys - Ade triple drop-out medium and subsequently grown on SC -Leu to maintain the pP9-V_H_H plasmid. Spot tests were realized on SC-His plates, grown at different temperatures during 3 days before scores were attributed. The score represents the number of dilutions on which a robust growth is visible with a V_H_H minus the number of dilutions on which a robust growth is visible with the empty plasmid containing no V_H_H (“pP9” on Figure 2B).

For GFP experiments, GFY216 to 219 and GFY225 cells were grown at 30°C overnight in SC-Leu, diluted to 9x10^6^ cells/mL in the morning and arrested in G1 by adding 1 μg/mL alpha factor. After 2 hours, cells were washed and released in fresh medium at 30°C. Alpha factor arrest and cell cycle restart was confirmed by microscope observation. Cells were collected at time 0, 120 minutes and 180 minutes after alpha factor release and analyzed on a MACSQuant Analyzer (Miltenyi Biotec). The fraction of GFP-positive cells was determined at each time point, using the FlowJo software (version 10.8.1). All flow cytometry results are shown in Supplemental Table S1.

### Production and purification of V_H_H proteins

For miniproduction of the four V_H_H (E02, F03, F08 and H01), HEK293 cells were plated in 6 well plates at 700 000 cells per well 2 days before transfection. Cells were transfected with 2.5 µg of Minibody DNA and 7.5 µL XtremeGene HP DNA. 24h post transfection the medium was replaced by medium without serum. After 6 days, the medium was collected, and the production was checked by Western blot with an anti-rabbit antibody coupled to HRP. The expected size for V_H_H-Fc is around 42 kDa. Production and purification of V_H_H H01 was performed at the Institut Pasteur protein production facility, as follows. The V_H_H H01 was expressed in Expi293F cells at 3717°C and 8% CO_2_ (Expi293, ThermoFisher Scientific). Cell culture supernatants were collected 4 days post transfection and purified by protein G chromatography (Cytiva). Washing step was done with DPBS 1X pH 7.2 and V_H_H were eluted with Glycine 0.1M pH 2.5 in wells containing TRIS 1M pH 8.0.

### Dot blot experiments with purified V _H_H proteins

Hybond-XL nylon membrane (Amersham-GE Healthcare) was soaked in 2X SSC for 20 minutes. Synthetic oligonucleotides, plasmidic or genomic DNA were denatured in 50-75 μL 0.4 M NaOH, incubated at 95°C for 2 minutes, and rapidly put on ice. They were transferred on the nylon membrane under limited vaccum, using a homemade dot blotter. For each sample, 500 μL of 0.4 M NaOH were added in each well, three times. After disassembly of the dot blotter, the membrane was incubated one hour in PBS. Blocking was performed for one hour in TBST (20 mM Tris pH7.5, 150 mM NaCl, 0.1% Tween 20) supplemented with 5% dry milk. V_H_H were diluted 1/50^th^ to 1/500^th^ in preliminary tests in TBST + 5% dry milk and incubated overnight at 4°C. Purified H01 V_H_H was diluted 1/1000^th^ in TBST + 5% dry milk in subsequent experiments. The secondary anti-rabbit antibody (Thermo Scientific) was diluted 1/2000^th^, incubated for one hour in TBST + 5% dry milk at room temperature, and revealed by ECL Prime Western Blotting (Amersham). Signals were revealed on a Bio-Rad Chemidoc and quantifications performed using the Image Lab software (version 5.2). For hybridization on yeast genomic DNAs, the quantification reflects the average of three independent experiments (Figure 3D).

### Secondary structure capture on HiTrap column

Plasmidic secondary structures were captured on an Äkta start FPLC (Cytiva) at a flow rate of 1 mL/min for all buffers, unless otherwise specified. A 1 mL HiTrap protein G (Cytiva) was first equilibrated with 10 mL 20 mM sodium phosphate buffer (pH 7.0). The V_H_H (20-500 μg, depending on the experiment) was loaded in 10 mL of the same buffer. To lower non-specific binding, the column was saturated with 15 mL 1% BSA in 20 mM sodium phosphate buffer. The pRW3222 plasmid (Fabre et al. 2002) was heat denatured for 10’ at 95°C before being loaded on the column in 5 mL T10N50 (10 mM Tris pH 8.0, 50 mM NaCl). In order to reduce heat degradation of the plasmid, 10 mM MgCl_2_ was added just before the heating step (Marguet and Forterre 1998). Right after plasmid injection, 1 mL fractions were collected until the end of the experiment (ca. 30 fractions). The column was washed with 10 mL of the same buffer. Elution of plasmidic DNA bound to the V_H_H was performed with 10 mL T10N200 (10 mM Tris pH 8.0, 200 mM NaCl). The column was finally washed with 5 mL of T10N50.

For each fraction, 100 μL were concentrated using a Concentrator plus (Eppendorf) and loaded on a 1% agarose gel. Transfer and hybridization with the radioactively labeled pUC19 plasmid were performed as previously described (Viterbo et al. 2018). Signal quantification was performed as previously (Viterbo et al. 2016).

## Acknowledgements

This work was partially supported by a grant from the AFM-Telethon to G.-F. R.. Work in G.-F. R. laboratory is generously supported by the Institut Pasteur and the CNRS.

---

**Supplemental Figure S1: Phage display enrichment. A:** Enrichment determined by the output/input ratio, after each round of selection. **B:** Enrichment determined by ELISA assay for the CTG hairpin and for the GC clamp.

**Supplemental Figure S2: pRW3222 binding to protein G column in the presence or absence of the V**_H_**H** The same amount of plasmid (20 μg) was loaded on a protein G column with or without prior binding of the V_H_H (flow rate= 1mL/min). The relaxed form as well as several supercoiled forms are visible after migration and ethidium bromide coloration. In the presence of the nanobody (upper panel), the plasmid is eluted when the salt concentration is increased to 200 mM Na^+^ and detected in fractions 15-18. In the absence of the V_H_H (lower panel), no plasmid elution is detected when the salt concentration was increased. Denat.: plasmid was denatured before loading on the gel. Non denat.: plasmid was not denatured before loading. T10N50: 10 mM Tris pH 8.0, 50 mM NaCl; T10N200: 10 mM Tris pH 8.0, 200 mM NaCl. NaP: 20 mM sodium-phosphate pH 7.0.

**Supplemental Figure S3: Visualization of DNA structures on digested and denatured pRW3222** The CTG-containing plasmid was linearized before heat denaturation and rapid renaturation. It was loaded on the protein G column and the high salt fraction was observed by electron microscopy. Single-stranded regions adopting different conformations, hairpin-like or more complex structures are indicated by a white arrow. There is one such region on each plasmid.

**Supplemental Figure S4: Visualization of DNA structures on native pRW3222** The CTG-containing plasmid was directly loaded on the protein G column and the high salt fraction was observed by electron microscopy. White arrows point to DNA structures (see text for details). Note that some plasmids seem to carry two structured regions.

## References

Ait Saada A, Costa AB, Sheng Z, et al (2021) Structural parameters of palindromic repeats determine the specificity of nuclease attack of secondary structures. Nucleic Acids Res 49:3932–3947. 10.1093/nar/gkab168

Arandel L, Espinoza MP, Matloka M, et al (2017) Immortalized human myotonic dystrophy muscle cell lines to assess therapeutic compounds. Dis Model Mech 10:487–497. 10.1242/dmm.027367

Axford MM, Wang YH, Nakamori M, et al (2013a) Detection of Slipped-DNAs at the Trinucleotide Repeats of the Myotonic Dystrophy Type I Disease Locus in Patient Tissues. PLoS Genet 9:1–13. 10.1371/journal.pgen.1003866

Axford MM, Wang Y-H, Nakamori M, et al (2013b) Detection of Slipped-DNAs at the Trinucleotide Repeats of the Myotonic Dystrophy Type I Disease Locus in Patient Tissues. PLOS Genet 9:e1003866. 10.1371/journal.pgen.1003866

Coïc E, Richard G-F, Haber JE (2006) Saccharomyces cerevisiae donor preference during mating-type switching is dependent on chromosome architecture and organization. Genetics 173:1197–1206. 10.1534/genetics.106.055392

Fabre E, Dujon B, Richard G (2002) Transcription and nuclear transport of CAG/CTG trinucleotide repeats in yeast. Nucleic Acids Res 30:3540–3547. 10.1093/nar/gkf483

Freudenreich CH, Kantrow SM, Zakian VA (1998) Expansion and length-dependent fragility of CTG repeats in yeast. Science 279:853–856

Freudenreich CH, Stavenhagen JB, Zakian VA (1997) Stability of a CTG/CAG trinucleotide repeat in yeast is dependent on its orientation in the genome. Mol Cell Biol 17:2090–2098

Gacy AM, Goellner G, Juranic N, et al (1995) Trinucleotide repeats that expand in human disease form hairpin structures in vitro. Cell 81:533–540

Kang S, Jaworski A, Ohshima K, Wells RD (1995) Expansion and deletion of CTG repeats from human disease genes are determined by the direction of replication in E. coli. Nat Genet 10:213–217

Leach DRF, Okely EA, Pinder DJ (1997) Repair by recombination of DNA containing a palindromic sequence. Mol Microbiol 26:597–606. 10.1046/j.1365-2958.1997.6071957.x

Liu G, Chen X, Bissler JJ, et al (2010) Replication-dependent instability at (CTG) x (CAG) repeat hairpins in human cells. Nat Chem Biol 6:652–9. 10.1038/nchembio.416

Lobachev KS, Gordenin DA, Resnick MA (2002) The Mre11 Complex Is Required for Repair of Hairpin-Capped Double-Strand Breaks and Prevention of Chromosome Rearrangements. Cell 108:183–193. 10.1016/S0092-8674(02)00614-1

Lobachev KS, Stenger JE, Kozyreva OG, et al (2000) Inverted Alu repeats unstable in yeast are excluded from the human genome. Embo J 19:3822–30

Marguet E, Forterre P (1998) Protection of DNA by salts against thermodegradation at temperatures typical for hyperthermophiles. Extremophiles 2:115–122. 10.1007/s007920050050

McMurray CT (1999) DNA secondary structure: a common and causative factor for expansion in human disease. Proc Natl Acad Sci USA 96:1823–1825

Mitas M (1997) Trinucleotide repeats associated with human diseases. Nucleic Acids Res 25:2245–2253

Mitas M, Yu A, Dill J, et al (1995a) Hairpin properties of single-stranded DNA containing a GC-rich triplet repeat: (CTG)15. Nucleic Acids Res 23:1050–1059

Mitas M, Yu A, Dill J, Haworth IS (1995b) The trinucleotide repeat sequence d(CGG)15 forms a heat-stable hairpin containing Gsyn.Ganti base pairs. Biochemistry 34:12803–12811

Moutel S, Bery N, Bernard V, et al (2016) NaLi-H1: A universal synthetic library of humanized nanobodies providing highly functional antibodies and intrabodies. eLife 5:e16228. 10.7554/eLife.16228

Nakamori M, Panigrahi GB, Lanni S, et al (2020) A slipped-CAG DNA-binding small molecule induces trinucleotide-repeat contractions in vivo. Nat Genet 52:146– 159. 10.1038/s41588-019-0575-8

Pearson CE, Tam M, Wang YH, et al (2002) Slipped-strand DNAs formed by long (CAG)*(CTG) repeats: slipped-out repeats and slip-out junctions. Nucleic Acids Res 30:4534–47

Pelletier R, Krasilnikova MM, Samadashwily GM, et al (2003) Replication and expansion of trinucleotide repeats in yeast. Mol Cell Biol 23:1349–1357. 10.1128/mcb.23.4.1349-1357.2003

Poggi L, Richard G-F (2021) Alternative DNA Structures In Vivo: Molecular Evidence and Remaining Questions. Microbiol Mol Biol Rev 85:. 10.1128/MMBR.00110-20

Richard G-F, Cyncynatus C, Dujon B (2003) Contractions and expansions of CAG/CTG trinucleotide repeats occur during ectopic gene conversion in yeast, by a MUS81-independent mechanism. J Mol Biol 326:769–782

Richard G-F, Dujon B, Haber JE (1999) Double-strand break repair can lead to high frequencies of deletions within short CAG/CTG trinucleotide repeats. Mol Gen Genet 261:871–882

Richard G-F, Goellner GM, McMurray CT, Haber JE (2000) Recombination-induced CAG trinucleotide repeat expansions in yeast involve the MRE11/RAD50/XRS2 complex. EMBO J 19:2381–2390

Samadashwily GM, Raca G, Mirkin SM (1997) Trinucleotide repeats affect DNA replication in vivo. Nat Genet 17:298–304. 10.1038/ng1197-298

Schweitzer JK, Livingston DM (1998) Expansions of CAG repeat tracts are frequent in a yeast mutant defective in Okazaki fragment maturation. Hum Mol Genet 7:69–74

Sheffield VC, Cox DR, Lerman LS, Myers RM (1989) Attachment of a 40-base-pair G + C-rich sequence (GC-clamp) to genomic DNA fragments by the polymerase chain reaction results in improved detection of single-base changes. Proc Natl Acad Sci 86:232–236. 10.1073/pnas.86.1.232

Steinmetzer K, Zannis-Hadjopoulos M, Price GB (1995) Anti-cruciform Monoclonal Antibody and Cruciform DNA Interaction. J Mol Biol 254:29–37. 10.1006/jmbi.1995.0596

Suen IS, Rhodes JN, Christy M, et al (1999) Structural properties of Friedreich’s ataxia d(GAA) repeats. Biochim Biophys Acta 1444:14–24

Viterbo D, Marchal A, Mosbach V, et al (2018) A fast, sensitive and cost-effective method for nucleic acid detection using non-radioactive probes. Biol Methods Protoc 3:. 10.1093/biomethods/bpy006

Viterbo D, Michoud G, Mosbach V, et al (2016) Replication stalling and heteroduplex formation within CAG/CTG trinucleotide repeats by mismatch repair. DNA Repair 42:94–106

Yu A, Barron MD, Romero RM, et al (1997) At physiological pH, d(CCG)15 forms a hairpin containing protonated cytosines and a distorted helix. Biochemistry 36:3687–3699

Yu A, Mitas M (1995) The purine-rich trinucleotide repeat sequences d(CAG)15 and d(GAC)15 form hairpins. Nucleic Acids Res 23:4055–4057

Zuker M (2003) Mfold web server for nucleic acid folding and hybridization prediction. Nucleic Acids Res 31:3406–3415. 10.1093/nar/gkg595

