## Supplementary material for "Imperfect hairpins formed by CTG trinucleotide repeats are heteroduplex DNA molecules in living cells": Table 1

Table 1: VHH clones identified by phage display

| **Clone name** | **Yeast plasmid** | **ELISA ratio** | **Redundancy** |
| --- | --- | --- | --- |
| A02 | pTRI136 | 61.6 | 7 |
| B02 | pTRI137 | 58.8 | 1 |
| C04 | pTRI138 | 42.3 | 2 |
| D11 | pTRI139 | 4.5 | 2 |
| E02 | pTRI140 | 36.7 | 5 |
| G05 | pTRI141 | 36.0 | 6 |
| H01 | pTRI142 | 65.1 | 15 |
| H10 | pTRI143 | 44.4 | 3 |
| C09 | pTRI144 | 55.1 | 22 |
| F08 | pTRI145 | 58.7 | 1 |
| F03 | pTRI146 | 7.5 | 4 |
