## Supplementary material for "Imperfect hairpins formed by CTG trinucleotide repeats are heteroduplex DNA molecules in living cells": Sup Fig S1

**A**

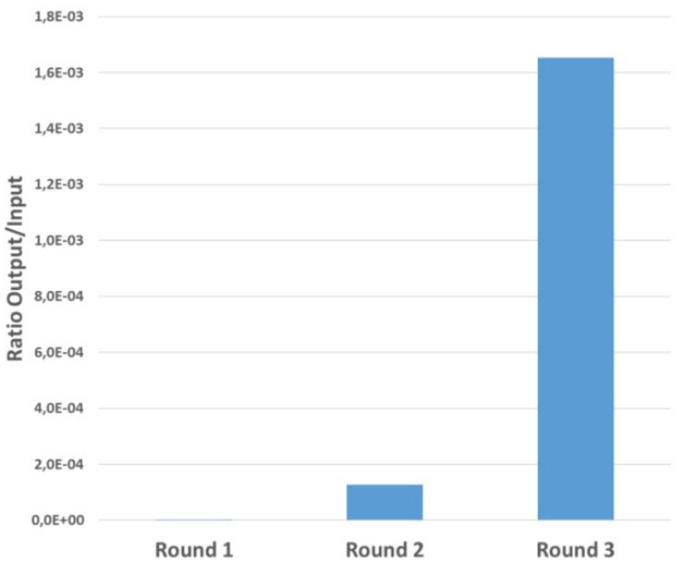


**Enrichment determined by output/input ratio**

| | Round 1 | Round 2 | Round 3 |
|---|---|---|---|
| Input | 8,1E+11 | 1,4E+12 | 1,3E+12 |
| Output | 7,2E+04 | 1,8E+08 | 2,1E+09 |
| Output/Input | 8,8E-08 | 1,3E-04 | 1,7E-03 |

**B**

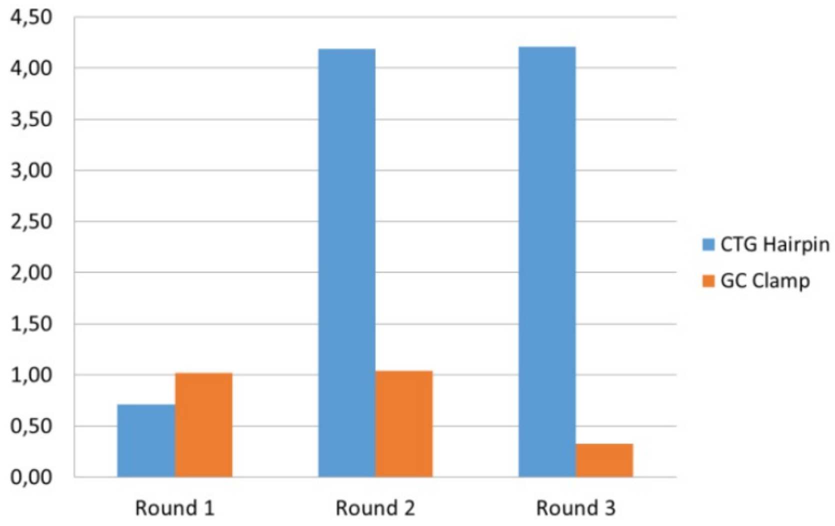


**Enrichment results by ELISA assay**
