## Supplementary figures and images for "Imperfect hairpins formed by CTG trinucleotide repeats are heteroduplex DNA molecules in living cells"

### Sup Fig S2

**HiTrap + $V_HH$**

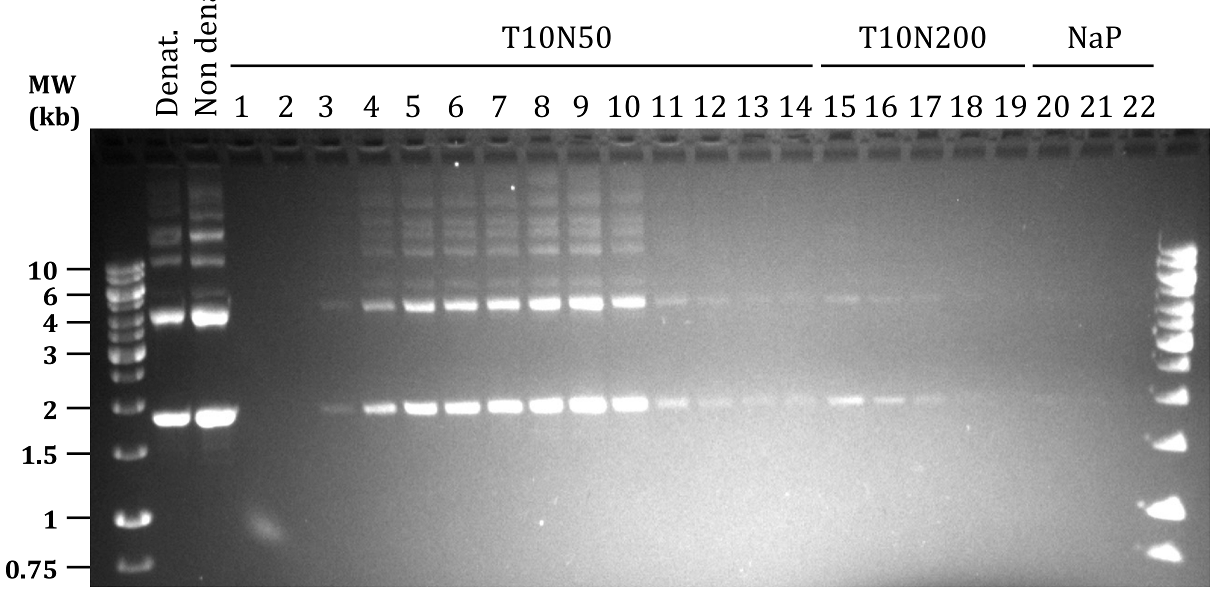


**HiTrap**

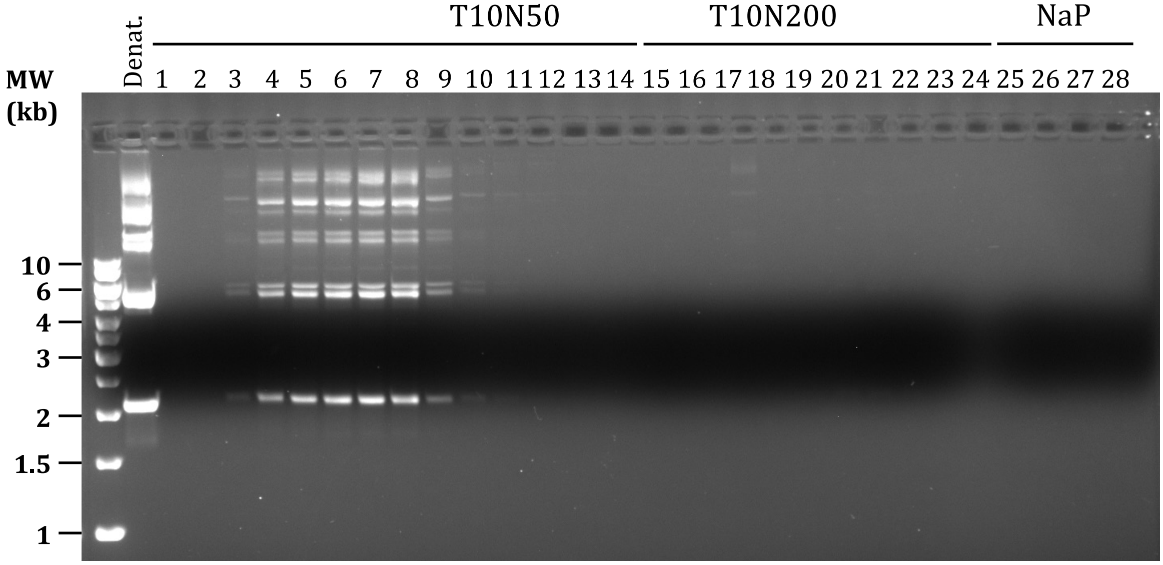

### Sup Fig S3

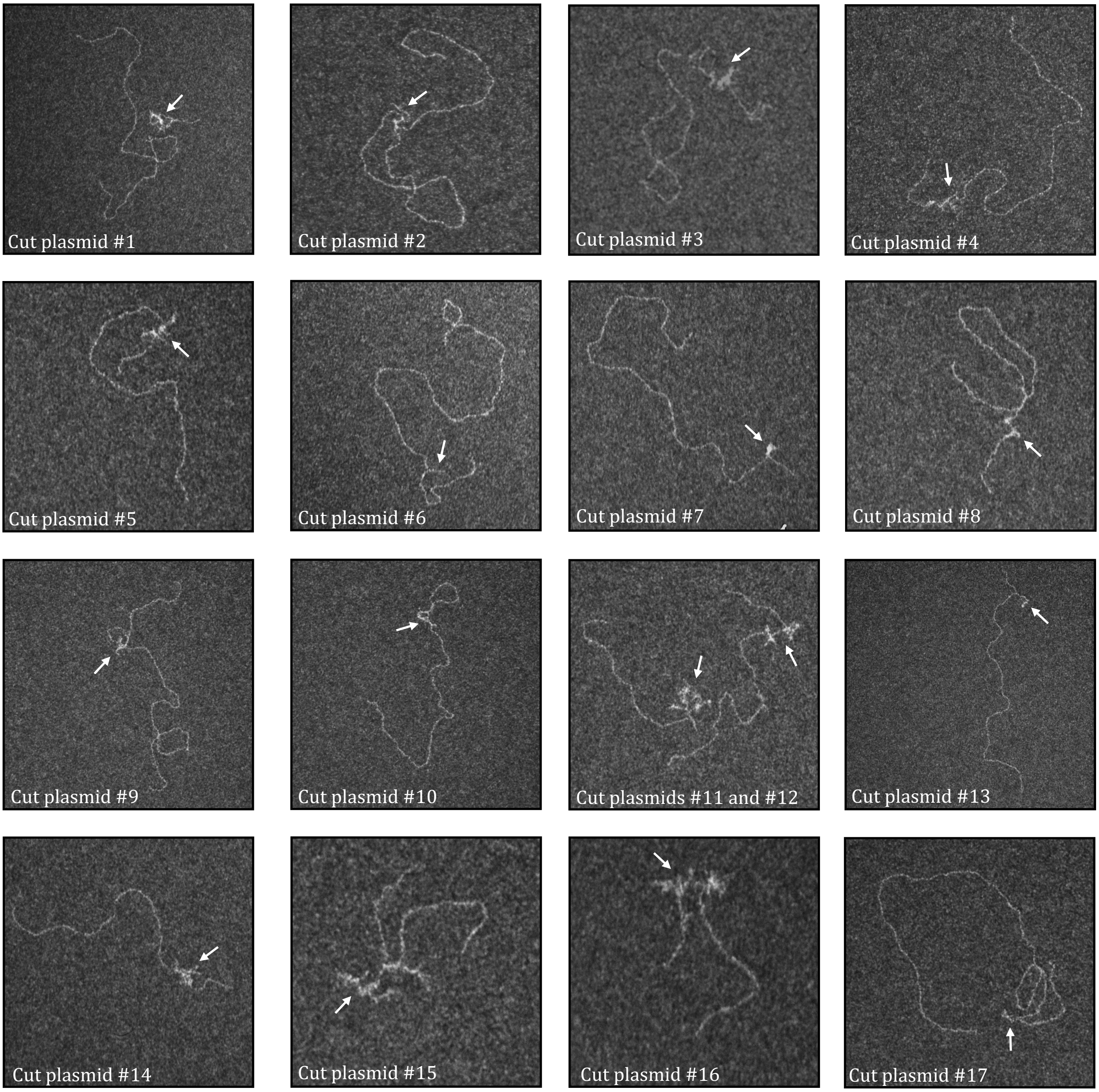

### Sup Fig S4

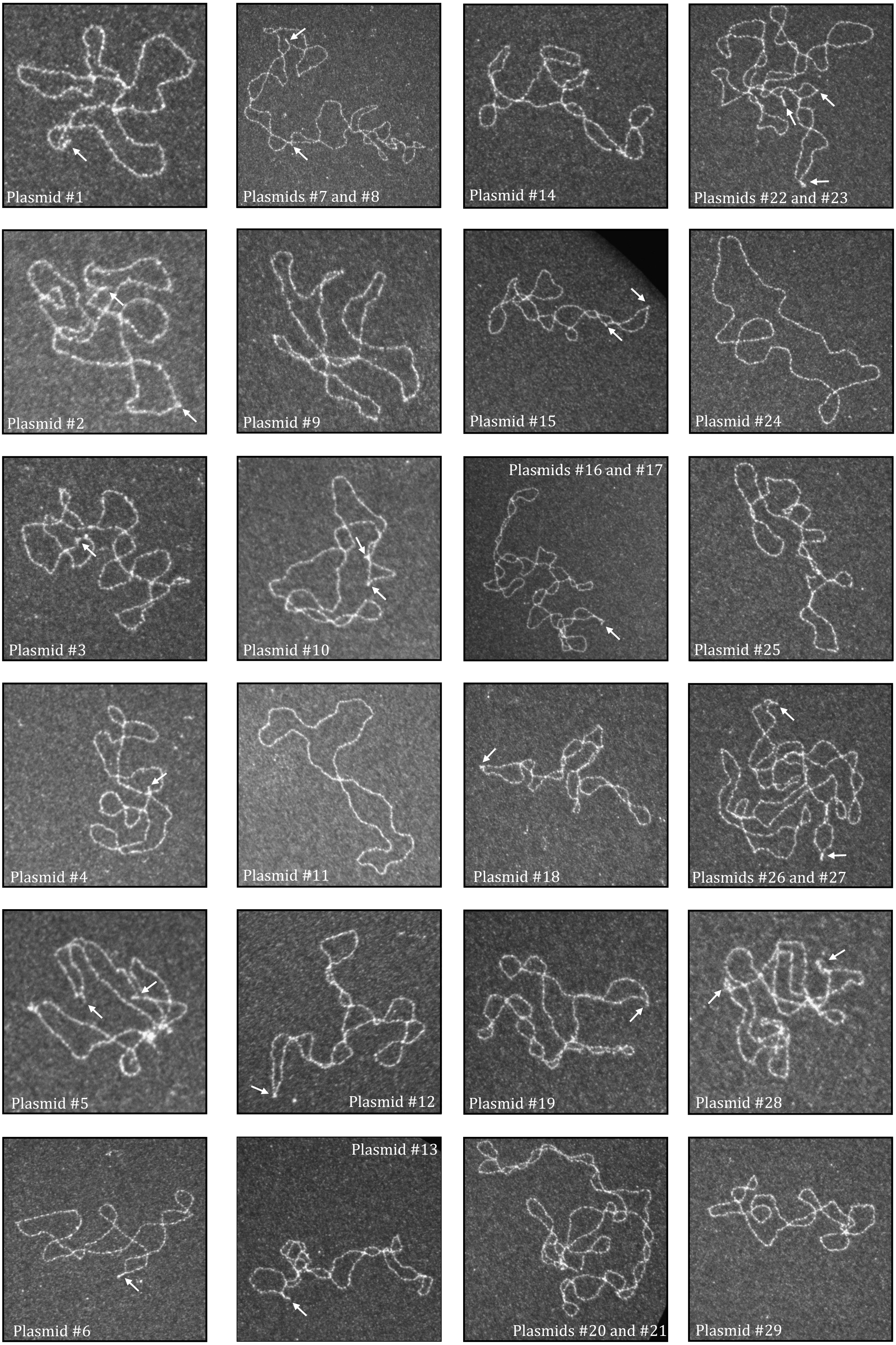

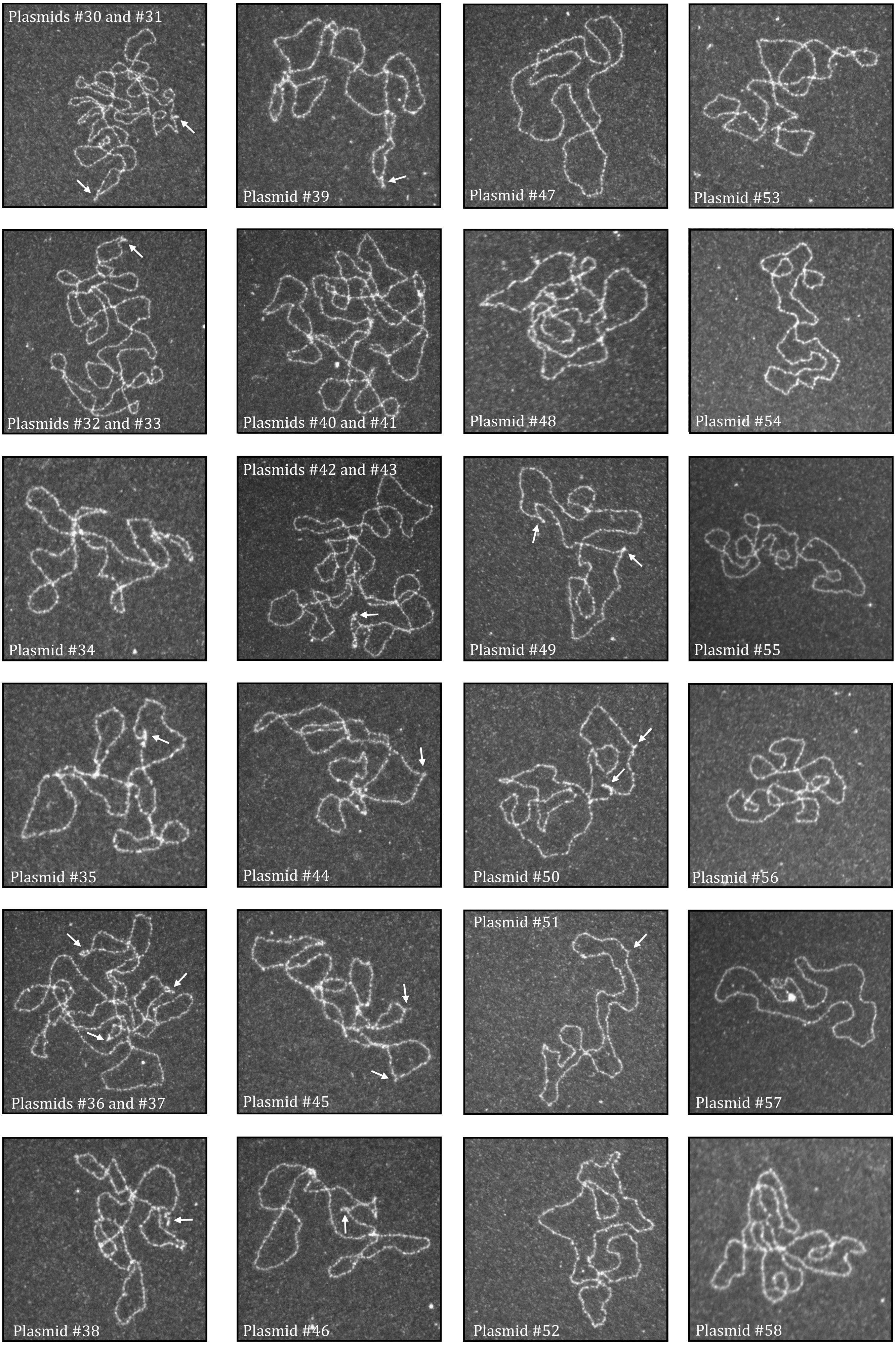
