## Supplementary material for "Imperfect hairpins formed by CTG trinucleotide repeats are heteroduplex DNA molecules in living cells": Sup Table S2

**Supplemental Table S2: List of primers used**

| **Name** | **Sequence** |
| --- | --- |
| VHHup* | **CCCACCAAACCCAAAAAAAGAGATCCTAGAACTAGCTATGGCCGGACGGGCCATGG**CGGAAGTGCAGCTGCAGGCTTC |
| VHHdown* | **ACCGGGCCTCTAGACACTAGCTACTCGAGGGGCCCCAGTGGCCCTATCTATGCGGCCGC**GCTACTCACAGTTAC |
| pB300-Sal | TCGACTCGAGTTAATTAACTAGTCTGCA |
| pB300-Pst | GACTAGTTAATTAACTCGAG |
| RAD27_1For | AATAATAGGGCAACGCGTCTGA |
| RAD27_2Rev | ATGTATCTATGCAAACGCCGAT |
| AR5 | ATTGCCTGTGCCAAGGGAATAT |
| repet_rad27 | GTTTATCTTGCCTGCTCATT |

* Homology with pP9 vector in bold
