## Supplementary material for "Imperfect hairpins formed by CTG trinucleotide repeats are heteroduplex DNA molecules in living cells": Sup Table S3

**Supplemental Table S3: yeast strains used in this study**

| **Name** | **Genotype** | **Origin** |
| --- | --- | --- |
| FYAT01 | *MAT*α *ura3*Δ851 *leu2*Δ1 *his3*Δ200 *trp1*Δ63 ade2-opal | Richard, Dujon and Haber, MGG 261: 871-882 (1999) |
| FYEF01-7C | *MAT*α *ura3*Δ851 *leu2*Δ1 *his3*Δ200 *trp1*Δ63 *ade2*-opal | FYAT01 |
| FYEF01-9C | *MAT*a *ura3*Δ851 *leu2*Δ1 *his3*Δ200 *lys2*Δ202 *ade2*-opal | FYAT01 |
| FYBL2-7B | *MAT*a *ura3*Δ851 *leu2*Δ1 *lys2*Δ202 *trp1*Δ63 | FYBL2 spore |
| GFY211 | *MAT*α *ura3*Δ851 *leu2*Δ1 *his3*Δ200 *trp1*Δ63 *ade2*-opal::(CTG)99 | FYEF01-7C |
| GFY212 | *MAT*α *ura3*Δ851 *leu2*Δ1 *his3*Δ200 *trp1*Δ63 *ade2*-opal *arg2*Δ::(CAG)30-eGFP | FYEF01-7C |
| GFY213 | *MAT*α *ura3*Δ851 *leu2*Δ1 *his3*Δ200 *trp1*Δ63 *ade2*-opal *arg2*Δ::(CTG)30-eGFP | FYEF01-7C |
| GFY214 | *MAT*α *ura3*Δ851 *leu2*Δ1 *his3*Δ200 *trp1*Δ63 *ade2*-opal *arg2*Δ::(GAA)30-eGFP | FYEF01-7C |
| GFY215 | *MAT*α *ura3*Δ851 *leu2*Δ1 *his3*Δ200 *trp1*Δ63 *ade2*-opal *arg2*Δ::(CGG)30-eGFP | FYEF01-7C |
| GFY216 | *MAT*a *ura3*Δ851 *leu2*Δ1 *his3*Δ200 *trp1*Δ63 *ade2*-opal *arg2*Δ::(CAG)30-eGFP | GFY212 |
| GFY217 | *MAT*a *ura3*Δ851 *leu2*Δ1 *his3*Δ200 *trp1*Δ63 *ade2*-opal *arg2*Δ::(CTG)30-eGFP | GFY213 |
| GFY218 | *MAT*a *ura3*Δ851 *leu2*Δ1 *his3*Δ200 *trp1*Δ63 *ade2*-opal *arg2*Δ::(GAA)30-eGFP | GFY214 |
| GFY219 | *MAT*a *ura3*Δ851 *leu2*Δ1 *his3*Δ200 *trp1*Δ63 *ade2*-opal *arg2*Δ::(CGG)30-eGFP | GFY215 |
| GFY225 | *MAT*a *ura3*Δ851 *leu2*Δ1 *lys2*Δ202 *trp1*Δ63 *arg2*Δ::(CTG)Δ-GFP | FYBL2-7B |
